# The C terminus of the mycobacterium ESX-1 secretion system substrate ESAT-6 is required for phagosomal membrane damage and virulence

**DOI:** 10.1101/2022.01.14.476355

**Authors:** Morwan M. Osman, Jonathan K. Shanahan, Frances Chu, Kevin K. Takaki, Malte L. Pinckert, Antonio J. Pagán, Roland Brosch, William H. Conrad, Lalita Ramakrishnan

## Abstract

*Mycobacterium tuberculosis* and its close relative *Mycobacterium marinum* infect macrophages and induce the formation of granulomas, organized macrophage-rich immune aggregates. These mycobacterial pathogens can accelerate and co-opt granuloma formation for their benefit, using the specialized secretion system ESX-1, a key virulence determinant. ESX-1-mediated virulence is attributed to the damage it causes to the membranes of macrophage phagosomal compartments, within which the bacteria reside. This phagosomal damage, in turn, has been attributed to the membranolytic activity of ESAT-6, the major secreted substrate of ESX-1. However, mutations that perturb ESAT- 6’s membranolytic activity often result in global impairment of ESX-1 secretion. This has precluded an understanding of the causal and mechanistic relationships between ESAT-6 membranolysis and ESX-1-mediated virulence. Here, we identify two conserved residues in the unstructured C-terminal tail of ESAT-6 required for phagosomal damage, granuloma formation and virulence. Importantly, these ESAT-6 mutants have near-normal levels of secretion, far higher than the minimal threshold we establish is needed for ESX-1-mediated virulence early in infection. Unexpectedly, these loss-of-function ESAT-6 mutants retain the ability to lyse acidified liposomes. Thus, ESAT-6’s virulence functions *in vivo* can be uncoupled from this *in vitro* surrogate assay. These uncoupling mutants highlight an enigmatic functional domain of ESAT-6 and provide key tools to investigate the mechanism of phagosomal damage and virulence.

**Significance Statement:** Tuberculosis (TB), an ancient disease of humanity, continues to be a major cause of worldwide death. The causative agent of TB, *Mycobacterium tuberculosis*, and its close pathogenic relative *Mycobacterium marinum*, initially infect, evade, and exploit macrophages, a major host defense against invading pathogens. Within macrophages, mycobacteria reside within host membrane-bound compartments called phagosomes.

Mycobacterium-induced damage of the phagosomal membranes is integral to pathogenesis, and this activity has been attributed the specialized mycobacterial secretion system ESX-1, and particularly to ESAT-6, its major secreted protein. Here, we show that the integrity of the unstructured ESAT-6 C-terminus is required for macrophage phagosomal damage, granuloma formation, and virulence.

## Introduction

The type VII secretion system ESX-1 (ESAT-6 Secretion System 1) is a major virulence determinant in *Mycobacterium tuberculosis* (Mtb) and its close relative *Mycobacterium marinum* (Mm) (1–4). ESX-1 was identified as a virulence determinant when a 9.4 kb deletion in this region was discovered in the live attenuated tuberculosis vaccine Bacillus Calmette–Guérin (BCG) (2, 5). This “Region of Difference 1 (RD1)” was found to be required for both Mtb and Mm virulence (1, 2). The *esx-1* locus was then found to extend beyond RD1 in both organisms (6, 7) (Fig. S1A). Mtb and Mm ESX-1 systems are functionally equivalent: complementation of Mm−ΔRD1 with a cosmid containing the Mtb ESX-1 locus restores ESX-1 functions (8), and the use of Mm has facilitated fundamental discoveries about ESX-1 function (9). ESX-1 promotes virulence through an array of processes including activation of cytosolic signaling pathways, macrophage death, and pathogenic acceleration of tuberculous granulomas through induction of MMP9, which in turn promote mycobacterial growth (3, 9–12). ESX-1 also mediates damage of the macrophage phagosomal membranes in which the bacteria reside, and this is thought to be integral to ESX-1-mediated virulence (13–15).

ESX-1’s membranolytic activity had been ascribed to its major secreted substrate ESAT-6 (6 kDa early secretory antigenic target) (16). ESAT-6 was discovered as a secreted, immunodominant Mtb antigen long before the *esx-1* locus was identified (17). Once the *esx-1* locus was identified, it was determined that the genes encoding ESAT-6 (*esxA*) and its secreted partner CFP-10 (*esxB*) reside in RD1 (Fig. S1A) (5, 6). ESAT-6 and CFP-10 (95 and 100 amino acids, respectively), were the first identified members of the type VII secretion-associated WXG100 superfamily, so named for their conserved WXG motif and their size of ∼100 amino acids (18). Pinning down the role of ESAT-6 and other ESX-1 substrates in membranolysis and virulence has been complicated by their co-dependency for secretion, as deletion of ESAT-6 causes loss of other ESX-1 substrates (19, 20). A separate line of evidence used purified recombinant ESAT-6 to implicate it in membranolysis. Purified ESAT-6, but not its co-secreted partner CFP-10, lysed artificial lipid bilayers, liposomes, and red blood cells (RBCs), leading to the conclusion that ESAT-6 functions as a classical pore-forming toxin (15, 21, 22).

However, we and others found that many of the pore-forming activities ascribed to ESAT-6, such as RBC lysis, was due to residual detergent contamination in ESAT-6 preparations made using standard, widely distributed protocols (8, 23, 24). Moreover, we found that true ESX-1-mediated RBC lysis was contact-dependent and caused gross membrane disruptions as opposed to distinct pores (8). These findings suggested that ESAT-6 was either not directly involved in ESX-1 mediated membranolysis *in vivo* or required additional mycobacterial factors. One such additional factor has been identified: we and others have shown that the mycobacterial cell surface lipid phthiocerol dimycocerosate (PDIM) is also required for macrophage phagosomal damage in both Mtb and Mm (25–29). There is also increasing evidence that ESX-1 substrates other than ESAT-6 are required for its pathogenic activity. Mm mutants in the ESX-1 genes *espE* and *espF* secrete normal levels of ESAT-6 but are attenuated in ESX-1-mediated virulence functions (30). These findings suggest that ESAT-6 is not sufficient for ESX-1 virulence function while leaving open the question of whether it is necessary. Continued efforts to understand to what extent ESAT-6 is directly involved in ESX-1 membranolysis and virulence have not provided clear answers. Mm transposon mutants have been identified that do not secrete ESAT-6 but are capable of damaging macrophage phagosomes as evidenced by bacterial cytosolic translocation (31), and ESAT-6 secretion-deficient ESX-1 mutants have been identified that are unaffected in intramacrophage growth and/or virulence in Mm and Mtb (32–36). These findings go against a direct role for ESAT-6 in ESX-1’s membranolytic activity. However, there is also strong evidence supporting ESAT-6’s direct involvement in ESX-1-mediated membranolytic activity and virulence. Levels of surface-bound ESAT-6 correlate with cytotoxicity and intramacrophage growth (35, 37). An N-terminal ESAT-6 point mutation (Q5K) preserves ESAT-6 secretion but attenuates phagosomal permeabilization, cytotoxicity and Mm growth in cultured macrophages, and zebrafish larvae (38).

Covalent modification of secreted ESAT-6 reduces hemolysis and intramacrophage growth (39), suggesting that ESAT-6 functions as a secreted effector. ESAT-6 Q5K and covalently modified ESAT-6 are both attenuated in ability to lyse acidified liposomes, which is considered a proxy for its *in vivo* membranolytic activity. Lysis of acidified liposomes is the single *in vitro* assay where both recombinant and natively purified Mtb ESAT-6 exhibit membranolytic activity in the absence of contaminating detergent (21, 23, 40).

In this work, we probe the role of ESAT-6 in virulence. We find that mutation of EccA1, a putative ESX-1 chaperone, or treatment with the drug ebselen results in a drastic reduction in ESAT-6 and of its co-secreted partner, CFP-10. Both the *eccA1* mutant and ebselen-treated Mm retain substantial phagosome-damaging activity, growth, and granuloma formation *in vivo*. In contrast, we find that two C-terminal point mutations in ESAT-6 allow substantial levels of ESAT-6 and CFP-10 secretion but cause complete loss of phagosomal membrane damage and virulence. Moreover, mutation of the C-terminus still allows for lysis of acidified liposomes, showing that there are additional requirements for ESAT-6 mediated membrane damage *in vivo*.

## Results

### Minimal ESAT-6 secretion suffices for ESX-1’s pathogenic functions

In prior work, we had found that a C-terminal M93T point mutation in ESAT-6 (Mm−ΔRD1::M93Tmt) resulted in greatly decreased secretion not only of ESAT-6, but also of CFP-10 (8). As CFP-10 is dependent on ESAT-6 for secretion, this suggested that the mutation might disrupt ESX-1 virulence simply by compromising ESAT-6-dependent ESX-1 substrate secretion (8). However, Mm mutants in *eccA1*, which encodes the AAA+ ATPase EccA1 – a putative chaperone for ESX-1 substrates – are also deficient for ESAT-6 and CFP-10 secretion (7, 33) yet, they are reported to be only somewhat compromised for virulence (33, 36). This suggested that minimal ESAT-6 secretion is compatible with ESX-1 mediated virulence. To study the link between ESAT-6 secretion and ESX-1-mediated virulence phenotypes, we examined a Mm transposon mutant in *eccA1* (Mm-*eccA1*::Tn). We confirmed that Mm-*eccA1*::Tn had minimal ESAT-6 and CFP-10 secretion, comparable to that of Mm−ΔRD1::M93Tmt (Fig. 1A and Fig. 3B) (8). We also confirmed that Mm-*eccA1*::Tn was deficient for RBC lysis, as previously reported (Fig. 1B) (7, 33). Next, we examined the extent to which Mm-*eccA1*::Tn could damage phagosomal membranes by using the galectin-8 staining assay. This assay takes advantage of the fact that cytosolic galectins bind to lumenal β-galactoside-containing glycans that become exposed on damaged vesicles and can be visualized by immunofluorescence microscopy (Fig. 1C) (28, 41, 42). Infection with Mm-*eccA1*::Tn resulted in decreased galectin-8 puncta but substantially more than an Mm-ΔRD1 (26% vs. 6.9% of wildtype puncta) (Fig. 1D). Also, as previously reported, Mm-*eccA1*::Tn had only partially attenuated growth in cultured macrophages (Fig. 1E) (7, 33). It achieved nearly wildtype bacterial burdens during zebrafish larval infection (0.82±0.16 fold of wildtype vs 0.29±0.05 fold for Mm−ΔRD1) and this was associated with wildtype levels of granuloma formation, an ESX-1-mediated phenotype (Fig. 1F-H) (3, 11, 12). Thus, the reduction in ESAT-6 secretion resulting in *eccA1*::Tn abrogates RBC lysis but retains significant phagosome-damaging activity and nearly wildtype levels of growth in human and zebrafish macrophages, at least over the first few days of infection.

**Figure 1.**
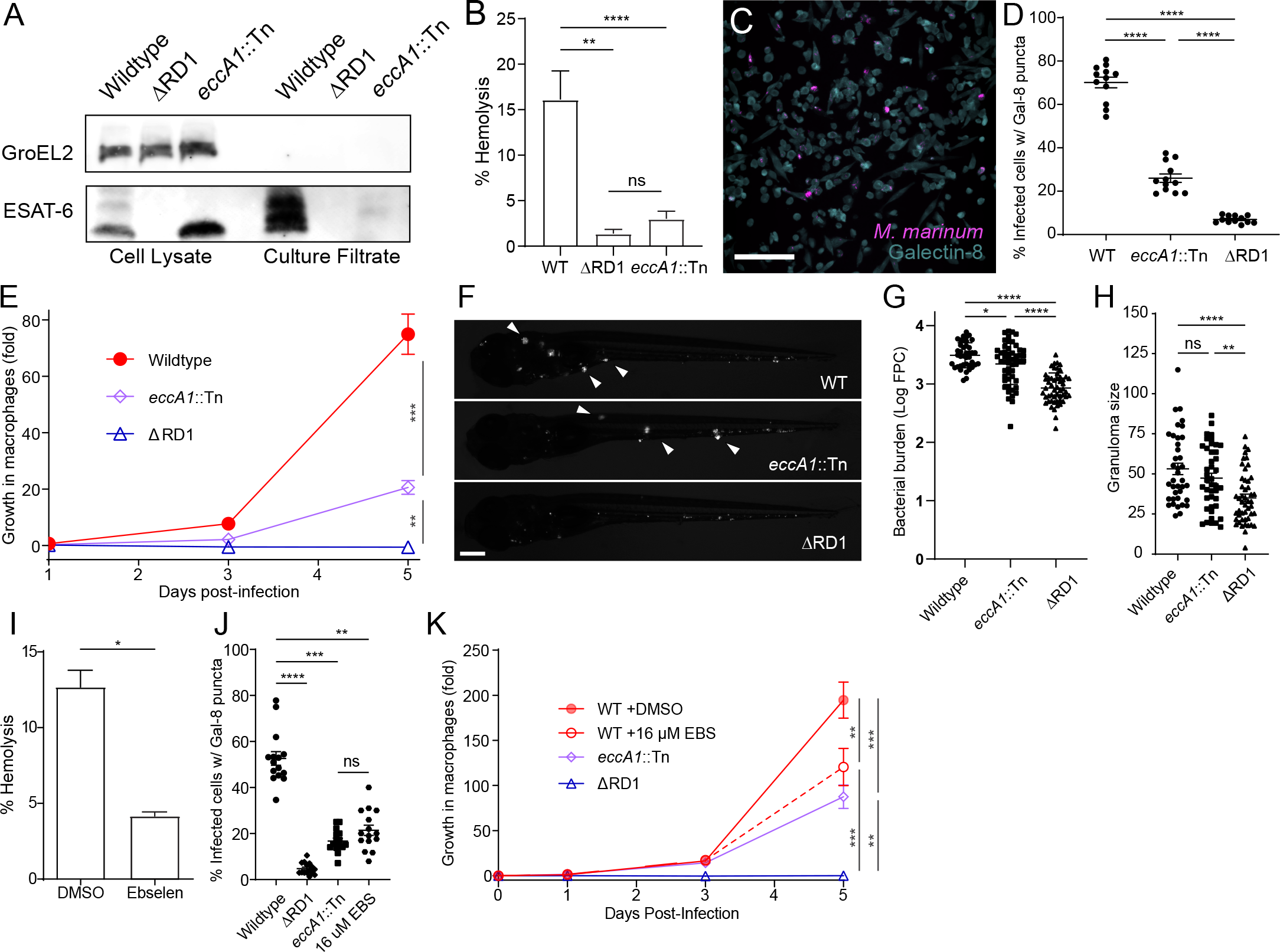
Minimal ESX-1 secretion is required for intramacrophage survival and virulence. (**A**) Immunoblot of 48-hour Mm cell lysates and culture filtrates. Data representative of three independent experiments. GroEL2 is shown as a lysis control. (**B**) Contact-dependent hemolysis of red blood cells by Mm. Data combined from four experimental replicates. (**C**) Representative image of THP-1 cells infected with Mm, stained for galectin-8. Scale bar, 100 μm. (**D**) Percent of infected THP-1 macrophages with galectin- 8 puncta. Each data point represents an individual imaging field. Horizontal lines, means. Statistics, one-way ANOVA with Šidák’s multiple comparisons test. (**E**) Intramacrophage growth of Mm within J774A.1 cells as measured by bacterial fluorescence. Data representative of three independent experiments. (**F**, **G**, **H**) Zebrafish larvae at 5 dpi. Data representative of four independent experiments. (**F**) Representative images. Scale bar, 250 μm. Arrowheads, granulomas. (**G**) Bacterial burdens as assessed by bacterial fluorescence. (**H**) Average infection foci size per larva. Statistics, one-way ANOVA with Dunnett’s test. (**I**) Contact-dependent hemolysis of red blood cells by Mm treated with vehicle or 16 μM Ebselen. Data combined from four experimental replicates. (**J**) Percent of infected THP-1 macrophages with galectin-8 puncta. Each data point represents an individual imaging field. Horizontal lines, means. Statistics, one-way ANOVA with Šidák’s multiple comparisons test. (**K**) Intramacrophage growth of J774A.1 cells infected with Mm. Data representative of three independent experiments. (**E**, **K**) One-way ANOVA with Bonferroni’s multiple comparisons test. Statistics, **** = p<0.0001, *** = p≤0.001 , ** = p<0.01, * = p<0.05, ns = p>0.05.

To further investigate the relationship between levels of ESAT-6 secretion, membranolysis and virulence, we used the drug ebselen. Ebselen reduces ESAT-6 secretion in wildtype Mm to similar levels as Mm-*eccA1*::Tn albeit through an EccA1- independent mechanism (43). Ebselen resulted in the same dissociation between ESAT-6 secretion and virulence phenotypes in wildtype Mm that we had observed for Mm- *eccA1*::Tn., with inhibition of RBC lysis and phagosomal damage combined with a minimal effect on intracellular bacterial growth (Fig. 1I-K). Zebrafish experiments with ebselen were precluded by drug toxicity at concentrations exceeding 1 μM, a dose at which ebselen shows no effect on ESX-1 function (43). In sum, minimal ESAT-6 secretion is sufficient to support some level of phagosomal damage, which, in turn, is sufficient for substantial mycobacterial virulence at least early in infection. Given this insight, the reduction in ESX-1 substrate secretion in Mm−ΔRD1::M93Tmt was not sufficient to explain its complete loss of phagosomal damage and virulence, warranting further investigation of the role of ESAT-6 and, particularly, its C-terminus.

### ESAT-6 C-terminal point mutants retain appreciable ESAT-6 secretion

Residues 83-95 of ESAT-6 form a highly conserved C-terminal motif (Fig. 2A-C, Fig. S3) (44). Our prior work on the effect of the ESAT-6 M93T mutation had used an RD1 deletion mutant of Mm complemented with a cosmid containing the extended ESX- 1 region of Mtb expressing an ESAT-6 M93T mutant (Δ*esxA*::M93IMtb) (Fig. 2A-C, Fig. S1). For a more refined analysis, we constructed an ESAT-6 deletion mutant (Mm−Δ*esxA*) and complemented it using an integrating plasmid with wildtype ESAT-6 or ESAT-6 M93T mutation (Fig. S1). We also generated a second mutant of ESAT-6 in this region, where another highly conserved methionine was changed to an isoleucine (Δ*esxA*::M83IMtb) (Fig. 2A-C). Like the ESAT-6 M93T mutant, the ESAT-6 M83I mutant also displayed reductions in secretion and intramacrophage growth in the context of cosmid complementation of Mm-ΔRD1 (Fig. S2) (8, 45).

**Figure 2.**
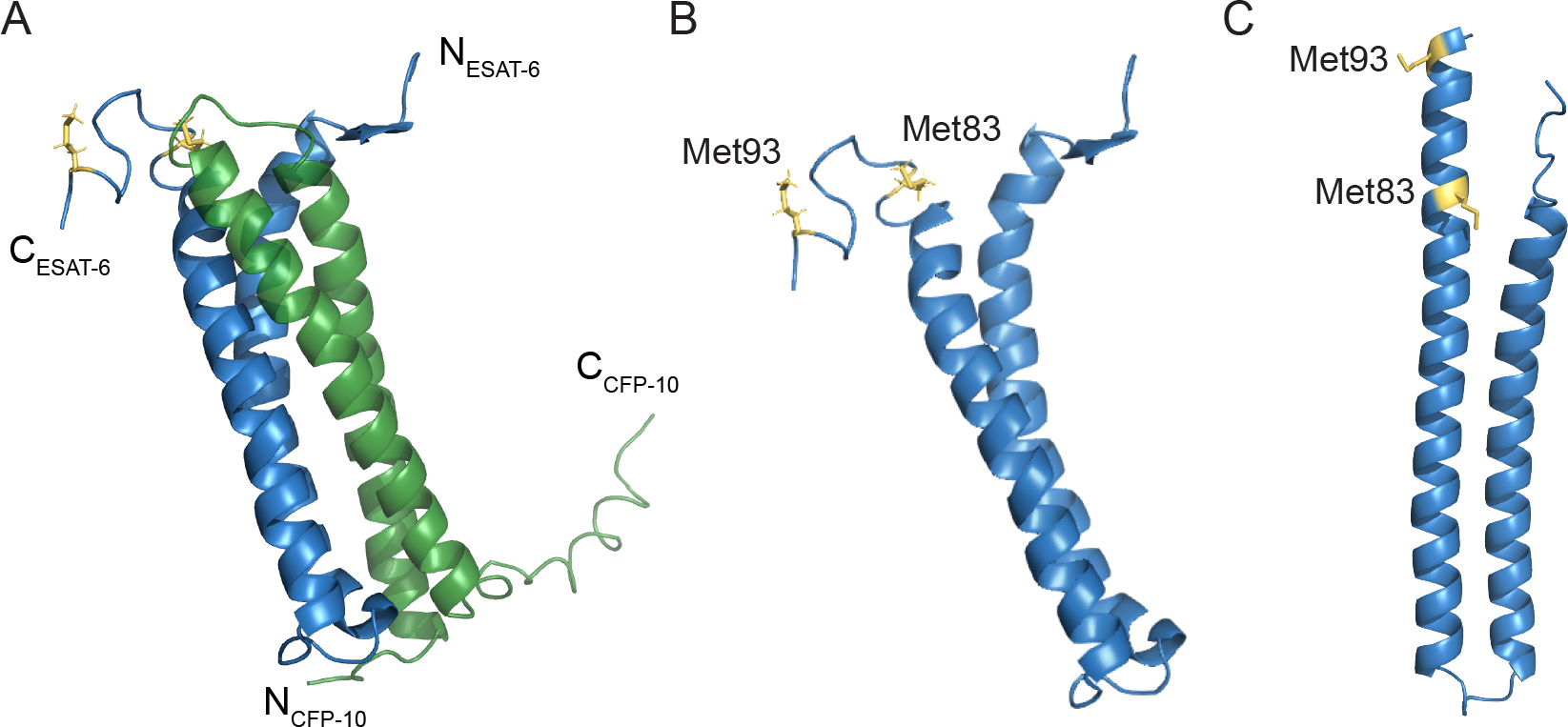
ESAT-6 structures with C-terminal point mutations. (**A**) NMR structure of the heterodimer formed by ESAT-6 (blue) and CFP-10 (green) (PDB 1WA8). N and C termini of ESAT-6 and CFP-10 are as labelled, as well as methionine 83 and 93 of ESAT-6 (yellow). (**B**, **C**) Structure of ESAT-6 alone with methionine residues 83 and 93 highlighted, as determined experimentally by NMR (**B**), or the predicted model using AlphaFold2 (**C**).

We found that Mm−Δ*esxA* had total loss of ESAT-6 and CFP-10 secretion, similar to Mm−ΔRD1, and this was restored by *esxA* complementation (Fig. 3A). Complementation of Mm−Δ*esxA* with ESAT6 M93T (Δ*esxA*::M93TMtb) restored secretion to a substantial degree, more than the original cosmid complementation system and much more than Mm-*eccA1*::Tn (Fig. 3A and B). Thus, the reduced secretion seen with the cosmid complementation of Mm−ΔRD1 is likely an artifact of partial complementation of the RD1 locus, gene dosage effects, subunit stoichiometry effects or a combination of these that are mitigated by the current system. Importantly, the two C-terminal mutations largely preserve ESAT-6 and CFP-10 secretion.

**Figure 3.**
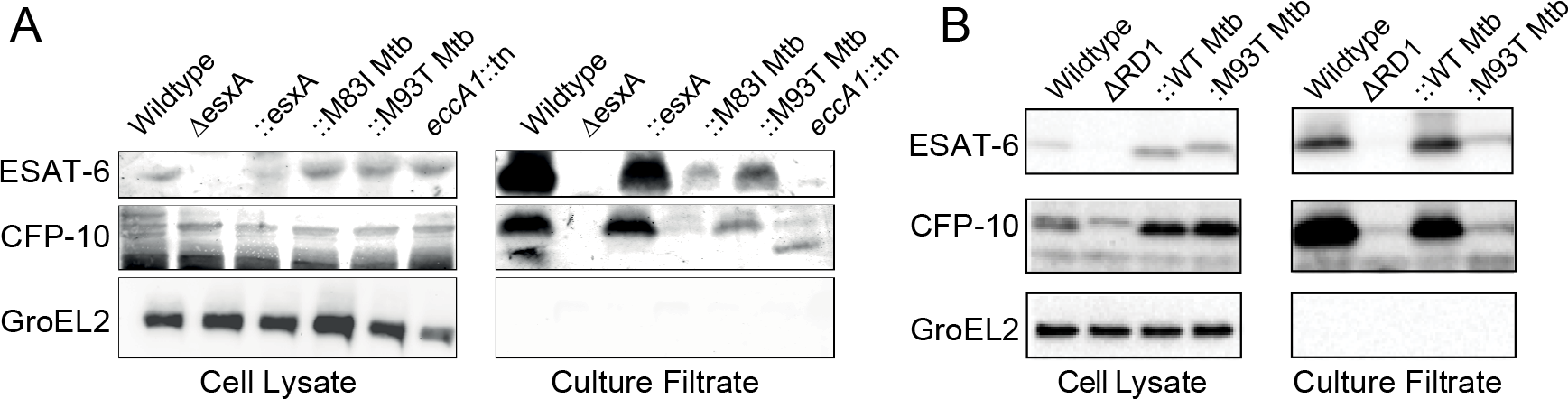
C-terminal point mutations in ESAT-6 support substantial levels of ESAT- 6 and CFP-10 secretion. (**A**, **B**) Immunoblots of 48-hour Mm cell lysates and culture filtrates. Representative of three independent experiments. (**B**) Reprinted with permission from (8).

### ESAT-6 C-terminal point mutants lose membranolytic activity and virulence

We could now use the two mutants to study the impact of ESAT-6 C-terminal integrity on ESAT-6 membranolytic activity and virulence. We first ascertained that Mm−Δ*esxA* was compromised for RBC lysis activity, phagosomal damaging activity, macrophage growth, and growth and granuloma formation in zebrafish larvae, similar to Mm−ΔRD1 (Fig. 4A-F) (8, 11, 28). All of these defects were rescued by complementation with Mtb ESAT-6 (Mm−Δ*esxA*::WTMtb) (Fig. 4A-F). However, ESAT- 6 bearing either C-terminal mutation failed to complement any of the phenotypes. Thus, the integrity of the ESAT-6 C terminus, while largely dispensable for secretion, is absolutely required for phagosomal membrane damage, which, in turn, is linked to granuloma formation and mycobacterial growth in vivo.

**Figure 4.**
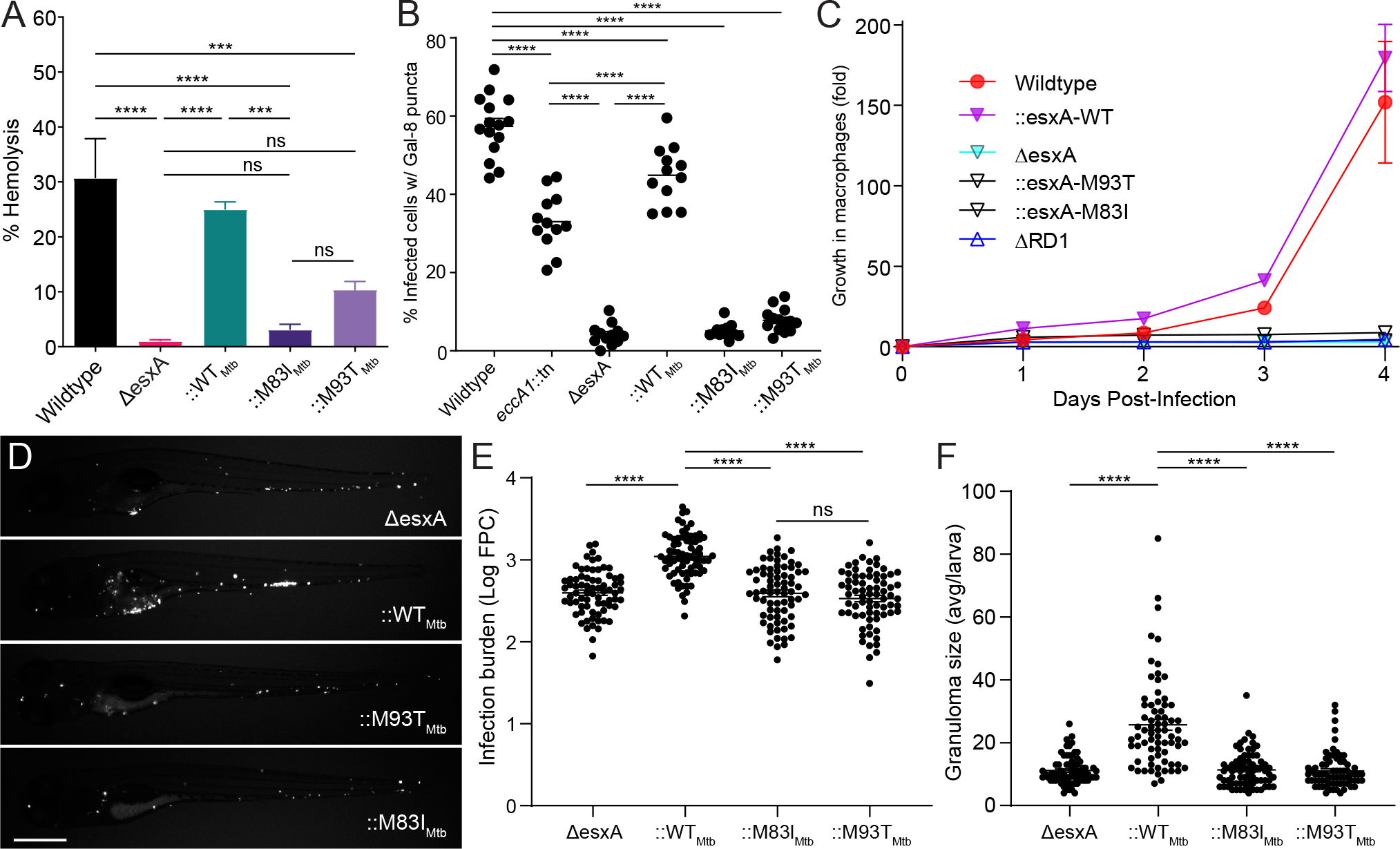
ESAT-6 mediates phagosomal damage and virulence. (**A**) Contact-dependent hemolysis of RBC by Mm. Data combined from four experimental replicates. (**B**) Percent of infected THP-1 macrophages with galectin-8 puncta. Each data point represents individual imaging fields. Horizontal lines, means. Statistics, one-way ANOVA with Šidák correction for multiple comparisons. (**C**) Intramacrophage growth of Mm within J774A.1 macrophages as measured by bacterial fluorescence. Data representative of four independent experiments. (**D**,**E**,**F**) Zebrafish larvae at 5 dpi. (**D**) Representative images. Scale bar, 500 μm. (**E**) Bacterial burdens as assessed by bacterial fluorescence. Statistics, one-way ANOVA with Dunnett’s test. (**F**) Average infection foci size per larva. Statistics, one-way ANOVA with Dunnett’s test. Data representative of three independent experiments. Statistics, **** = p<0.0001, *** = p≤0.001, ns = p>0.05.

### The ESAT-6 M93T mutant retains wildtype levels of acidified liposome lysis activity

Under acidic conditions (≤pH 5), ESAT-6 undergoes a conformational shift and inserts into liposomal membranes, resulting in their lysis (16, 46). Based on these observations, ESAT-6’s lysis of acidified liposomes has been used as a proxy for its role in phagosomal damage and permeabilization (16, 23). However, the relevance of the liposome lysis assay for ESAT-6’s role in phagosomal damage has been unclear. During Mm and Mtb infection, inhibition of phagosomal acidification, using the vacuolar type ATPase (v-ATPase) inhibitors bafilomycin A1 or concanamycin A, either has no effect on or even enhances phagosomal permeabilization/damage (8, 26, 47, 48). However, it has been argued that these findings may represent artifacts caused by alterations in phagosomal membrane composition caused by chemical inhibition of v-ATPase (49). To address the role of phagosomal acidification under natural conditions, we took advantage of the observation that during macrophage infection, mycobacteria reside in both acidified and nonacidified phagosomes (50). We found that 24 hours post-infection, 88.5% of Mm (415/469), were in acidified phagosomes as determined by staining with LysoTracker, an acidophilic dye that labels lysosomal compartments (50). Co-staining with galectin-8 showed that a significantly higher proportion of bacteria residing in acidified phagosomes were positive (2-fold increase over nonacidified) (Fig. 5). To see if phagosomal acidification also increases Mtb-mediated phagosomal damage, we used the auxotrophic Mtb H37Rv strain, mc^2^6206, a containment level 2 organism, which retains ESX-1 function (43, 51, 52). After 24 hours, 67.4% (320/475) of the bacterial phagosomes were acidified. As with Mm, there was significant increase in galectin-8 recruitment to bacteria residing in acidified compartments (1.7-fold increase over nonacidified) (Fig. 5). These findings show that both Mm and Mtb ESAT-6-mediated phagosomal damage is indeed enhanced by acidification.

**Figure 5.**
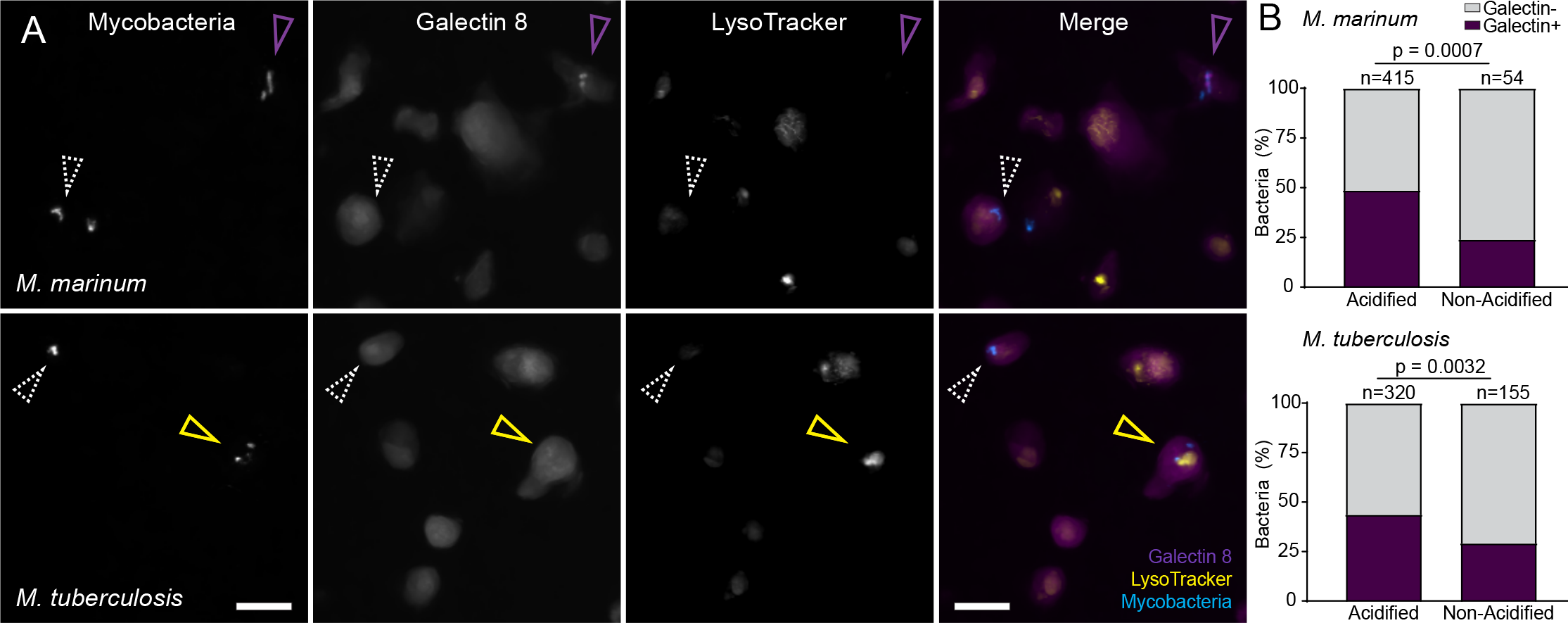
Mm- and Mtb-mediated phagosomal membrane damage is enhanced in acidified phagosomes. (**A**,**B**) Galectin-8 labeled, LysoTracker Red-stained THP-1 cells at 24 hours post-infection with EBFP2-expressing Mm (top) or Mtb (bottom). (**A**) Representative images of mycobacteria, Galectin-8, or Lysotracker, shown individually or as a merge of all three channels. Scale bar, 10 μm. Magenta arrowhead, bacteria proximal to galectin-8 puncta. Yellow arrowhead, acidified bacteria proximal to galectin-8 puncta. White arrowhead, bacteria that did not induce galectin-8 puncta. (**B**) Percent of bacteria located proximal to galectin-8 puncta, sorted by colocalization with LysoTracker-positive compartments. Statistics, Fisher’s exact test.

Importantly, these findings re-enforced the argument that the lytic activity of ESAT-6 on acidified liposomes is a proxy of ESAT-6-mediated phagosomal damage (16). Our finding that the C-terminal mutants had almost no phagosomal damage (Fig. 4B), when most of the bacteria are in acidified phagosomes, indicated that they are unable to damage phagosomes even under the sensitizing acidified condition. Therefore, we expected that the C-terminal mutations would cause ESAT-6 to lose acidified liposome lysis activity. We used recombinant ESAT-6-M93T, a less conservative mutation than M83I, which we hypothesized to be more likely to disrupt ESAT-6 function. To measure lysis, we generated 1,2-Dioleoyl-sn-glycero-3-phosphocholine (DOPC) liposomes containing the fluorescent dye 8-Aminonaphthalene-1,3,6-Trisulfonic acid (ANTS), a formulation which has been used for previous systematic analyses of ESAT-6’s lytic activity (46, 53). We were surprised to find that ESAT-6-M93T retained wildtype levels of acidified lysis (Fig. 6). This stood in contrast to the prior findings that recombinant ESAT-6 containing the N-terminal Q5K mutation, which caused loss in phagosomal damage in Mm, was found to be deficient in lysis of acidified DOPC liposomes (38). Thus, our finding uncouples ESAT-6’s *in vitro* and *in vivo* membrane-damaging activities even under acidified conditions, suggesting an *in vivo* role for specific residues in the C terminus.

**Figure 6.**
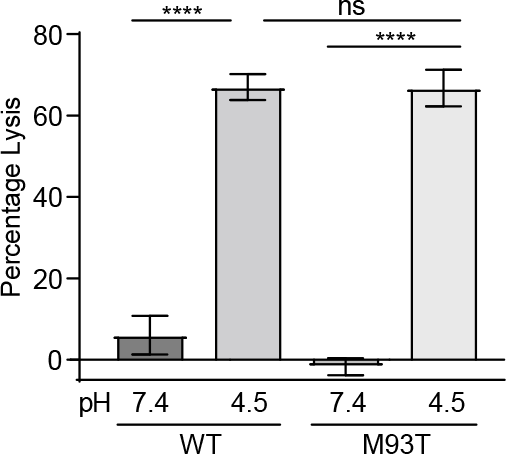
Recombinant ESAT-6-M93T has wild-type levels of acidified liposome lysis activity. Quantification of pH-dependent liposome lysis by 3 µM recombinant ESAT-6 as measured by fluorescent ANTS release from DOPC liposomes. Statistics, one-way ANOVA with Šidák’s multiple comparisons test; **** = p<0.0001, ns = p>0.05. Data combined from four experimental replicates.

## Discussion

This work was instigated by our finding that barely detectable levels of ESAT-6 secretion in both an Mm *eccA1* mutant and ebselen treated Mm only halved ESX-1- mediated phagosomal membrane. In turn, this level of phagosomal damage was sufficient for relatively high levels of *in vivo* growth compared to ESX-1 deficient Mm. Our findings led us to revisit our previous conclusions that an ESAT-6 C-terminal point mutation (ESAT-6 M93T) lost phagosomal membrane damage and early virulence solely due to reduced secretion (8). The Mm-ESAT-6 M93T mutant allowed for much greater levels of ESAT-6 secretion than the Mm-*eccA1*::Tn mutant, showing that ESAT-6 C-terminal integrity was needed for ESAT-6’s direct membrane damaging and virulence effects. We solidified this conclusion through experiments demonstrating that the Mm- ESAT-6 M83I mutant had a similar dissociation between secretion and virulence phenotypes. In the course of this analysis, we found that *in vivo* ESAT-6-mediated phagosomal lysis by Mtb and Mm was enhanced in acidic compartments. This suggested that the pH-dependent lytic activity of purified ESAT-6 was a direct correlate of phagosomal damage. However, we found that purified ESAT-6-M93T retained full activity in lysing acidified liposomes.

Previous studies examining both transposon mutants and total knockouts of *eccA1* have found defects in hemolysis and *in vivo* growth similar to those we observed in Mm- *eccA1*::Tn (7, 36). A genome-wide Mtb transposon mutant screen in both cultured macrophages and the mouse TB model identified *eccA1*::Tn mutants as being attenuated (54–56). These findings are broadly consistent with ours, although in the absence of direct comparisons, it is not clear whether EccA1 is less important than the full ESX-1 locus in Mtb infection as well. Our findings that minimal ESAT-6 secretion is compatible with substantial preservation of early Mm infection phenotypes are also in line with prior observations of phagosomal damage (as reflected by cytosolic translocation of mycobacteria) in ESAT-6 secretion-deficient Mm mutants (31). We see two possible explanations for this. First, minute amounts of secretion that are not detected by immunoblotting could be sufficient for ESX-1 function (32, 57). Second, surface-bound ESAT-6 can partially compensate for the lack of secretion during early infection. Two studies have found that the retention of surface-bound ESAT-6 correlates with pathogenic phenotypes (35, 37). Furthermore, an earlier report on Δ*eccA1* found wildtype levels of ESAT-6 in bacterial cell surface extracts despite an overall reduction in ESX-1 secretion (36).

ESAT-6 binds to CFP-10 to form a heterodimeric complex and the two proteins are likely co-secreted (58). ESAT-6’s C-terminus does not participate in binding to CFP- 10 (Fig. 2A) and truncation of this region (Δ84-95) does not affect secretion, likely as CFP-10’s C-terminal tail is responsible for recognition and secretion of the heterodimer (58–61). However, conservation of ESAT-6’s C-terminus among mycobacterial homologs (Fig. S3), suggests it mediates an essential function beyond secretion. Consistent with this, an Mtb mutant expressing the C-terminal ESAT-6 truncation has reduced phagosomal permeabilization as reflected by translocation into the cytosol (59).

Furthermore, it was reported that complementation of *M. bovis* BCG with an ESX-1 cosmid expressing truncated ESAT-6 did not rescue attenuation (45). This region appears to be highly conserved in clinical strains: we examined the GMTV database of 2,819 Mtb isolates for occurrences of nonsynonymous (coding) mutations in *esxA*, and of the 8 occurrences of nonsynonymous mutations in *esxA*, none were in located in the conserved C-terminal motif (residues 83-95) (62).

Our finding that phagosomal acidification greatly enhances ESAT-6-mediated damage supports the model that ESAT-6’s pH-dependent functions *in vitro* are physiologically relevant (16, 21, 40). This assay does not fully capture ESAT-6’s *in vivo* role, as mycobacteria in non-acidified compartments still induced phagosomal damage to a greater extent than either of the C-terminal ESAT-6 mutants. However, it is likely that ESAT-6’s activity under acidic pH mimics an aspect of its *in vivo* function. This is further supported by the recent finding that membrane vesicles prepared using lipids from the THP-1 human monocytic cell line are also damaged by ESAT-6 (recombinant and native) only at acidic pH (23, 26). On this backdrop, it is interesting to consider the role of the ESAT-6 C terminus in phagosomal damage, particularly of the two point mutants examined here. It has been shown that in the context of acidified liposome lysis, the ESAT-6 C-terminus does not insert into the membrane and remains solvent exposed (46). NMR and crystal structures of the heterodimer predict that the tail is floppy and unstructured (44, 58). However, the authors who solved the crystal structure suggested that this region could also adopt an alpha helical structure *in vivo* (44). This was based on identification of a C-terminal motif conserved across actinobacterial ESAT-6 homologs that is consistent with an alpha helical structure (44). This hypothesis is supported by the recent AlphaFold2 structural prediction of ESAT-6 with a structured C terminus (63) (Fig. 2C). This disparity between experimental and predicted structure could reflect a conformational versatility of the C-terminus that is required for different aspects of ESAT-6’s *in vivo* functions.

The ESAT-6 C-terminal tail resides on the exterior of lipid membranes following insertion (46). While an ESAT-6 C-terminal truncation mutant can insert into liposome membranes, it has reduced lytic activity (46). This suggests a direct role for the C terminus in enabling “basal” ESAT-6 membranolytic function. The ESAT-6 M93T and M83I point mutants are relatively conservative. As methionine and isoleucine are both small hydrophobic residues, an isoleucine mutation is highly conservative, while threonine, as a small polar residue would be less so. Neither of these mutations would be predicted to be particularly disruptive to C-terminal structure, and in this light, perhaps it is not surprising after all that the M93T mutation does not affect its *in vitro* basal membranolytic activity. Rather, this finding suggests that the M93T mutant is defective in a C-terminally mediated function that is required for robust *in vivo* lysis. Following membrane insertion, the C-terminus could enhance oligomerization of ESAT-6, interactions with mycobacterial or host factors, or act as a bacterial sensor for contact-dependent lysis by ESX-1, or any combination of these. Putative mycobacterial factors include other ESX-1 substrates that contribute to virulence in addition to ESAT-6, such as EspA, EspE, or EspF (9, 32, 64), or oligomers of EspB and EspC which are proposed to be part of a putative extracellular secretory complex mediating contact-dependent lysis (65, 66). A systematic mutational analysis of the C terminus coupled with biochemical studies to identify mycobacterial, or possibly even host, determinants that bind to wildtype but not M93T and other *in vivo* deficient ESAT-6 mutants may pave the way to a fuller understanding of ESAT-6’s *in vivo* function.

## Materials and Methods

### Bacterial strains and culture methods

All Mm and Mtb strains used are listed in Table S3. Mm strains were all derived from wildtype Mm purchased from American Type Culture Collection (strain M, ATCC #BAA-535). Mm strains were grown without agitation at 33°C in 7H9 Middlebrook’s medium (Difco) supplemented with 10% OADS and 0.05% Tween®80 (Sigma) with appropriate selection [50 μg /mL hygromycin (Mediatech) and 20 μg /mL kanamycin (Sigma)] or on supplemented Middlebrook 7H10 agar (Millipore) [no Tween®80] (67). Mtb mc^2^6206 strains (51) were grown without agitation at 37°C in 7H9 Middlebrook’s medium (Difco) supplemented with 10% OADC (Becton Dickinson), 0.05% Tween®80 (Sigma), 12 μg/mL pantothenic acid (Sigma) and 25 μg /mL L-leucine (Sigma) with appropriate selection [20 μg /mL kanamycin (Sigma)] or on supplemented Middlebrook 7H10 agar (Millipore) [no Tween®80].

### Generation of mutant strains

The *eccA1*::Tn mutant was isolated from a Mm transposon mutant library. Briefly, a novel *mariner*-based mini transposon containing an excisable hygromycin-resistance cassette was used to mutagenize wildtype Mm in small batches and individual colonies were isolated and sequenced. The transposon insertion in *eccA1*::Tn was confirmed by semi random, nested PCR and was located 16.26% into the *eccA1* open reading frame, as measured by distance from the ATG start site.

The Mm-Δ*esxA* mutant was generated using phage transduction (68). Briefly, a fragment containing sequences flanking *esxA* (23-955 bp upstream of the ATG, 13-1101 bp and downstream of its stop codon) were cloned into pYUB854 to generate pYUB854-*esxA*. The resulting cosmid was digested with PacI (NEB) and the insert was ligated into phAE159 (a gift from William Jacobs) to generate the phSP105 phasmid. Following ligation, packaging, and transduction, phSP105 was transformed into *M. smegmatis* mc^2^155. Harvested phages were transduced into wildtype Mm and plated on 7H10-Hyg. Positive colonies were confirmed by southern blotting, and the hygromycin cassette was excised via transformation with pYUB870. Loss of pYUB870 was confirmed by plating on 7H10+sucrose plates, and excision of cassette confirmed via PCR.

### Complementation Constructs

Mm-Δ*esxA* was complemented with pMH406 (69). ESAT-6-M83I and ESAT-6-M93T complementation constructs (Table S1) were generated via site directed mutagenesis using primers listed in Table S2.

### Single-cell suspensions of mycobacteria

Single-cell suspensions of mycobacteria were prepared as previously described (67), with minor modifications. Briefly, mycobacteria were cultured with antibiotic selection to mid-log (OD600 = 0.3 – 0.6) and then pelleted at 3,220 x g for 20 minutes at 20°C. The top layer of the pellet containing precipitated kanamycin was discarded following gentle scraping with a disposable inoculating loop. The underlying pellet was resuspended in 5 mL 7H9 media supplemented with 10% OADS (no Tween®80) and then passed 10 times through a 27-gauge blunt needle. Mycobacteria were then pelleted at 100 x g for 1 minute. A total of 4 mL supernatant was collected. The bacterial pellet was resuspended in 5 mL of 7H9 OADS. Mycobacteria were again passed 10 times through a blunt needle, pelleted at 100 x g, and supernatant collected. This was repeated until 15 mL supernatant was collected. The supernatant was then passed through a 5 μm syringe filter (Pall, 4650) to isolate single-cell bacteria. Single-cell bacteria were concentrated by centrifugation at 3,220 x g for 30 minutes at room temperature. The pellet was then resuspended in 200 μL 7H9 OADS, divided into 5 μL aliquots, and stored at −80°C. Mycobacterial concentration was determined by plating for CFU.

### Intramacrophage growth assays

Growth assays were performed as described previously with minor modifications (8). J774A.1 cells were maintained in Gibco Dulbecco’s Modified Eagle Medium (DMEM). 24 hours before infection, cells were scraped and resuspended in DMEM to a concentration of 2.5 x 10^5^ cells/mL. J774A.1s were plated in a 24-well optical bottom tissue culture plate (Perkin Elmer, 1450-606) by aliquoting 500 μL of this cell suspension to each well and then incubating at 37°C overnight. The following day, cells were washed twice with PBS and infected with antibiotic-free media containing single-cell suspensions of tdTomato-expressing Mm at a multiplicity of infection of ∼0.25 (wildtype, *eccA1*::Tn, ::*esxA*) or ∼0.5 (attenuated strains) for 4 hours at 33°C, 5% CO2. After infection, cells were washed twice with PBS, 500 μL fresh media added to each well and then incubated at 33°C, 5% CO2. Cells were imaged by fluorescence microscopy at indicated timepoints using a Nikon Eclipse Ti-E equipped with a Ti-S-E Motor XY Stage, a C-HGFIE 130-W mercury light source, a 23/0.10 Plan Apochromat objective, and a Chroma ET-CY3 (49004) filter cube. Fluorescence images were captured with a Photometrics CoolSNAP HQ2 Monochrome Camera, using NIS-Elements (version 3.22). Resulting images were analyzed in ImageJ using a custom script for fluorescence pixel count (FPC), to determine bacterial burden (67).

### Galectin-8 immunofluorescence

THP-1 cells were diluted to 5 x 10^5^ cells/mL in RPMI + 10% FCS (Gibco) and treated with 100 nM phorbol 12-myrystate-13-acetate (PMA) (Sigma, P1585). THP-1 cells were plated in a 24-well optical bottom tissue culture plate (Perkin Elmer, 1450-606) by aliquoting 500 μL of this cell suspension to each well and then incubating at 37°C for two days. PMA-containing media was then removed and replaced with fresh media, and the cells were allowed to rest for a day. The following day, cells were washed twice with PBS and infected with antibiotic-free media containing single-cell suspensions of tdTomato-expressing Mm at a multiplicity of infection of ∼1 for 4 hours at 33°C, 5% CO2. After infection, cells were washed twice with PBS, 500 μL fresh media added to each well and then incubated overnight at 33°C, 5% CO2.

Galectin-8 staining was done as previously described (28). Briefly, 24 hours after infection, cells were fixed in 4% (wt/vol) paraformaldehyde in PBS at room temperature for at least 30 minutes. Fixed cells were washed twice with PBS and then incubated in permeabilization/block (PB) buffer for 30 minutes at room temperature, and then stained with goat anti-human galectin-8 antibody (R&D Systems, AF1305) diluted in PB solution overnight at 4°C. Cells were then washed three times with PBS and stained with AlexaFluor488-conjugated donkey anti-goat IgG (ThermoFisher, A-11055) diluted in PB solution for one hour at room temperature. Cells were then washed three times in PBS and imaged.

LysoTracker experiments were conducted as above with the following modifications: cells were infected with EBFP2-expressing Mm or Mtb at an MOI of ∼0.5. The next day, cells were stained with 100 nM of LysoTracker Red DND-99 (Invitrogen) in RPMI for 45 minutes at 33°C, and then immediately fixed and stained as above. Acidic organelles, fluorescent bacteria, and galectin-8 puncta were identified using the 3D surface rendering feature of Imaris (Bitplane Scientific Software). Bacteria were scored as galectin-8 positive if they were located within 2 μm of a galectin-8 surface and as acidified if they were located within a LysoTracker surface.

### Zebrafish Husbandry

All zebrafish husbandry and experiments were performed in compliance with the UK Home Office and the Animal Ethics Committee of the University of Cambridge.

Zebrafish maintenance and spawning was performed as previously described (67). Briefly, zebrafish were maintained on a recirculating aquaculture system with a 14 hour light – 10 hour dark cycle. Fish were fed dry food and brine shrimp twice a day. Adult wildtype AB zebrafish were spawned, embryos collected and then housed in fish water (reverse osmosis water containing 0.18 g/L Instant Ocean) at 28.5°C. The fish water was supplemented with 0.25 μg/ml methylene blue from collection to 1 day post-fertilization (dpf), and at 1 dpf 0.003% PTU (1-phenyl-2-thiourea, Sigma) was added to prevent pigmentation.

### Zebrafish Infections

Zebrafish larvae were infected with the indicated strains of tdTomato-expressing Mm by injection into the caudal vein at 2 dpf using freshly thawed single-cell suspensions diluted to the same CFU / μL. Injection dose was plated on 7H10 + 10% OADS with appropriate antibiotic to ensure similar infection dose.

### Measuring Bacterial Burden in Larvae

Fluorescence microscopy was performed as previously described (67). Briefly, infected larvae were imaged by fluorescence microscopy at 5 days post-infection using a Nikon Eclipse Ti-E equipped with a Ti-S-E Motor XY Stage, a C-HGFIE 130-W mercury light source, a 23/0.10 Plan Apochromat objective, and a Chroma ET-CY3 (49004) filter cube. Fluorescence images were captured with a Photometrics CoolSNAP HQ2 Monochrome Camera, using NIS-Elements (version 3.22). The resulting images were analyzed in ImageJ using a custom script for fluorescence pixel count (FPC), to determine bacterial burden and infection foci size.

### Secretion Assay

Culture filtrate (CF) fractions and cell pellet (CP) fractions were prepared as described previously (8). Mm was grown to mid to late log stage, washed with PBS, resuspended to a final OD600 of 0.8-1.0 in 50 mL of modified Sauton’s Media, and incubated for 48 hours at 33°C. 10 μg of CP and 20 μg of CF were loaded per well for SDS-PAGE, and presence of ESAT-6, CFP-10 and GroEL2 were determined by western blotting with mouse anti-ESAT-6 clone 11G4 (1:3000; Thermo Fisher, HYB-076-08-02), rabbit anti-CFP-10 (1:5000; BEI, product NR13801), or mouse anti-GroEL2 clone IT-56 (1:1000; BEI, product NR-13655) primary antibody followed by HRP-conjugated goat anti-mouse (1:10,000; Stratech 115- 035-003-JIR) (Fig. 3B and S2), goat anti-mouse IgG DyLight™ 800 (1:15,000 Cell Signaling Technology #5257S), or goat anti-rabbit IgG DyLight™ 800 (1:15,000 Cell Signaling Technology #5151S) secondary antibody.

Chemiluminescence detection was then performed using Amersham ECL Western Blotting Detection Reagents (GE Lifesciences) and fluorescence imaging performed on a LI-COR Odyssey.

### Hemolysis Assay

Hemolytic activity was assessed as described previously (8). Briefly, 100 µL of 1% (v/v) sheep red blood cells (RBC) (Fisher Scientific) in PBS was combined with 100 µL of PBS, bacterial suspension or 0.1% Triton X-100 (Sigma), pelleted, and then incubated for two hours at 33°C. Pellets were resuspended, re-pelleted and the absorbance of the supernatant measured at 405 nm. Raw absorbance data were converted to percentage lysis, by subtracting PBS lysis (background lysis) and dividing by 0.1% Triton X-100 lysis (complete lysis).

### Protein Expression & Purification

Expression of recombinant ESAT-6 was conducted as previously (8) with the following modifications: Bacterial pellets were resuspended in 5 mL of Lysis Buffer (150 mM NaCl, 20 mM Tris pH 8, 5 mM Imidazole) per gram of pellet, and then sonicated on ice. Lysates were clarified by centrifugation at 15,000 x g for 30 minutes at 4°C. Clarified lysate was added to 1 mL of washed Ni-NTA bead slurry (Qiagen) and incubated for 1 hour at 4°C. Beads were then pelleted and washed with 10 column volumes (CVs) of lysis buffer. Next, beads were washed with 10 CV of binding buffer (150 mM NaCl, 20 mM Tris pH 8, 20 mM Imidazole), followed by 15 CV of wash buffer (150 mM NaCl, 20 mM Tris pH 8, 75 mM Imidazole). 3 CV of elution buffer (150 mM NaCl, 20 mM Tris pH 8, 1 M Imidazole) was added to elute protein. Eluate was concentrated and then loaded on a S75 size exclusion column (GE Healthcare) equilibrated with 150 mM NaCl, 20 mM Tris pH 8. Fractions were collected and monitored for recombinant protein elution and purity by SDS-PAGE followed by Coomassie staining. Protein concentration from highly purified eluted fractions were quantified by absorbance at 280 nm.

Constructs for mutant protein purification (Table S1) were generated via site directed mutagenesis using listed primers (Table S2).

### Liposome Lysis Assay

Liposomes containing 8-Aminonaphthalene-1,3,6-Trisulfonic Acid (ANTS) were generated as previously described with modifications (8). Briefly, 1,2-dioleoyl-sn- glycero-3-phosphocholine (Sigma) was dissolved in 86 μL of chloroform (Sigma) to 100 mg/mL. The resulting solution was transferred to a glass vial and chloroform evaporated under N2 gas and subsequently under vacuum overnight. Lipid films were resuspended in 400 μL of 125 mM ANTS (Invitrogen) in PBS by vortexing. Liposomes were generated by five freeze-thaw cycles by freezing in liquid nitrogen for three minutes and then thawing in a 40-50°C water bath for three minutes. Liposomes were then extruded twenty-five times through two 200 nm polycarbonate Nuclepore^TM^ filters (Whatman) and then washed in PBS by centrifugation at 60,000 × g for 30 minutes at 4°C. Liposomes were resuspended in 400 μL PBS for use in assays.

Stock solutions of 200 mM sodium phosphate and 100 mM citric acid with 150 mM NaCl were used to generate pH 4.5 and 7.4 phosphate-citrate pH buffers (pH buffer). The pH value of each buffer was confirmed after preparation. To measure liposome lytic activity of recombinant proteins, 5 μL of liposomes were resuspended in 200 μL pH buffer and 3 μM rESAT-6 or an equivalent volume of vehicle was added. Protein and liposome samples incubated with rotation at 20°C for 30 minutes in ultracentrifuge tubes. Intact liposomes were pelleted by centrifugation at 180,0000 × g for 30 minutes at 4°C and then ANTS fluorescence in the supernatants was measured [excitation/emission (nm): 350 ± 15/520 ± 15]. Raw fluorescence data were converted to percentage of lysis, by subtracting buffer only lysis (background lysis) and dividing by 0.05% Triton X-100 lysis (complete lysis) at each pH.

## Acknowledgments

We thank R. Hegde for insights and discussion and J. Blaza for support and encouragement. The following reagents were obtained through the NIH Biodefense and Emerging Infections Research Resources Repository, NIAID, NIH: monoclonal anti- Mycobacterium tuberculosis GroEL2 (gene Rv0440), clone CS-44 (produced in vitro), NR-13813. *M. tuberculosis* mc^2^6206 was a gift from William Jacob, pMH406 was a gift from David Sherman.

## Supplementary Information

### Supplementary Figures

**Figure S1.**
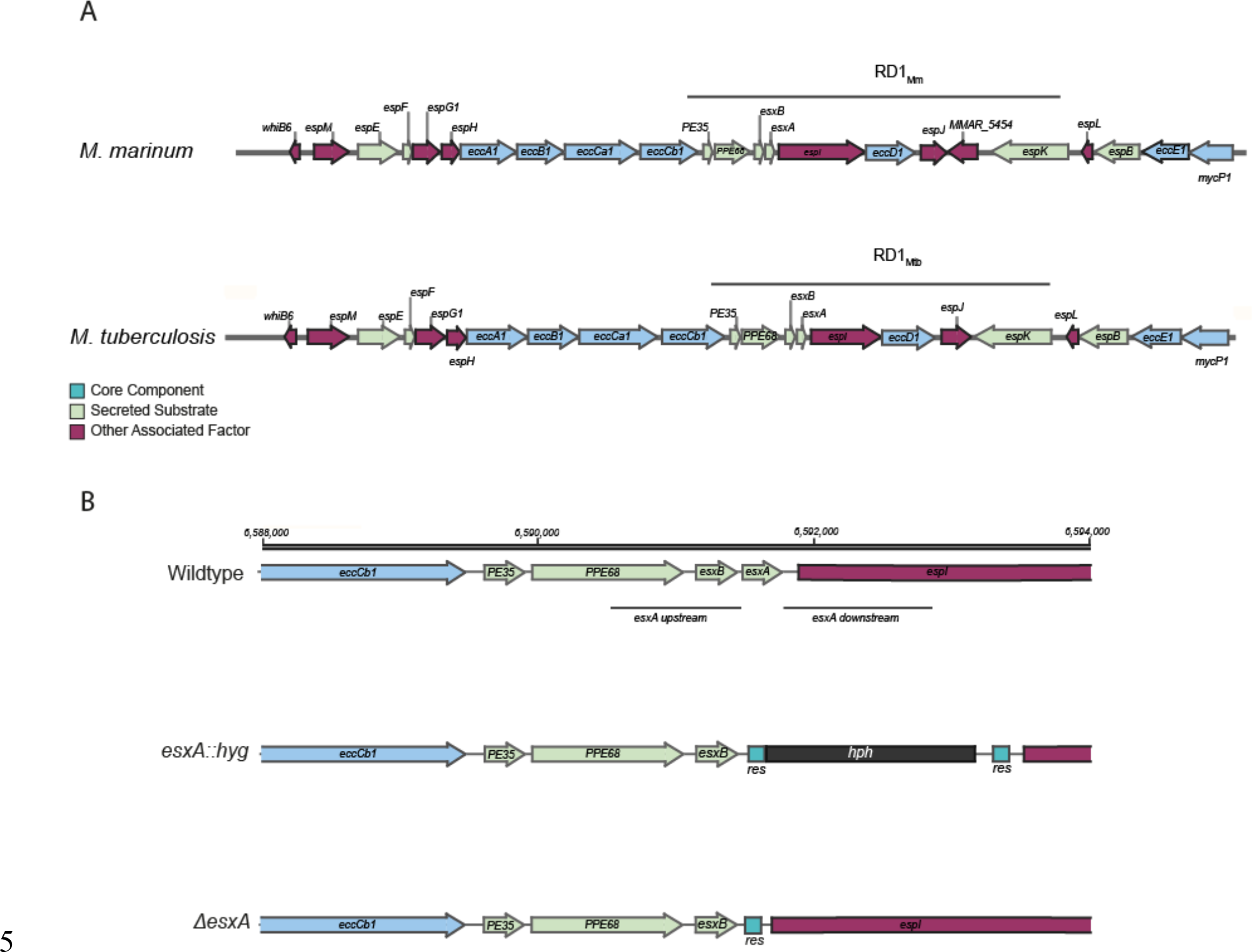
ESX-1 loci and scheme for Δ*esxA* mutant generation. (A) ) Alignment of Mm and Mtb ESX-1 loci, with regions corresponding to RD1 deletions. (**B**) Schematic showing the initial, intermediate, and final alleles in the generation of the *esxA* mutant in Mm. (Top) Wildtype *esxA* loci with flanking region upstream and downstream *esxA* as targeted by the deletion construct. (Middle) Phage transduction was used to generate the *esxA::hyg* mutant with *esxA* replaced by the *res*-flanked *hph* gene encoding the hygromycin-B-phosphotransferase selectable marker. (Bottom) The *hph* gene was then excised by a gamma-delta resolvase, generating the unmarked Mm-*ΔesxA* mutant. Full details of the primers, plasmids and phasmids can be found in the Materials & Methods.

**Figure S2.**
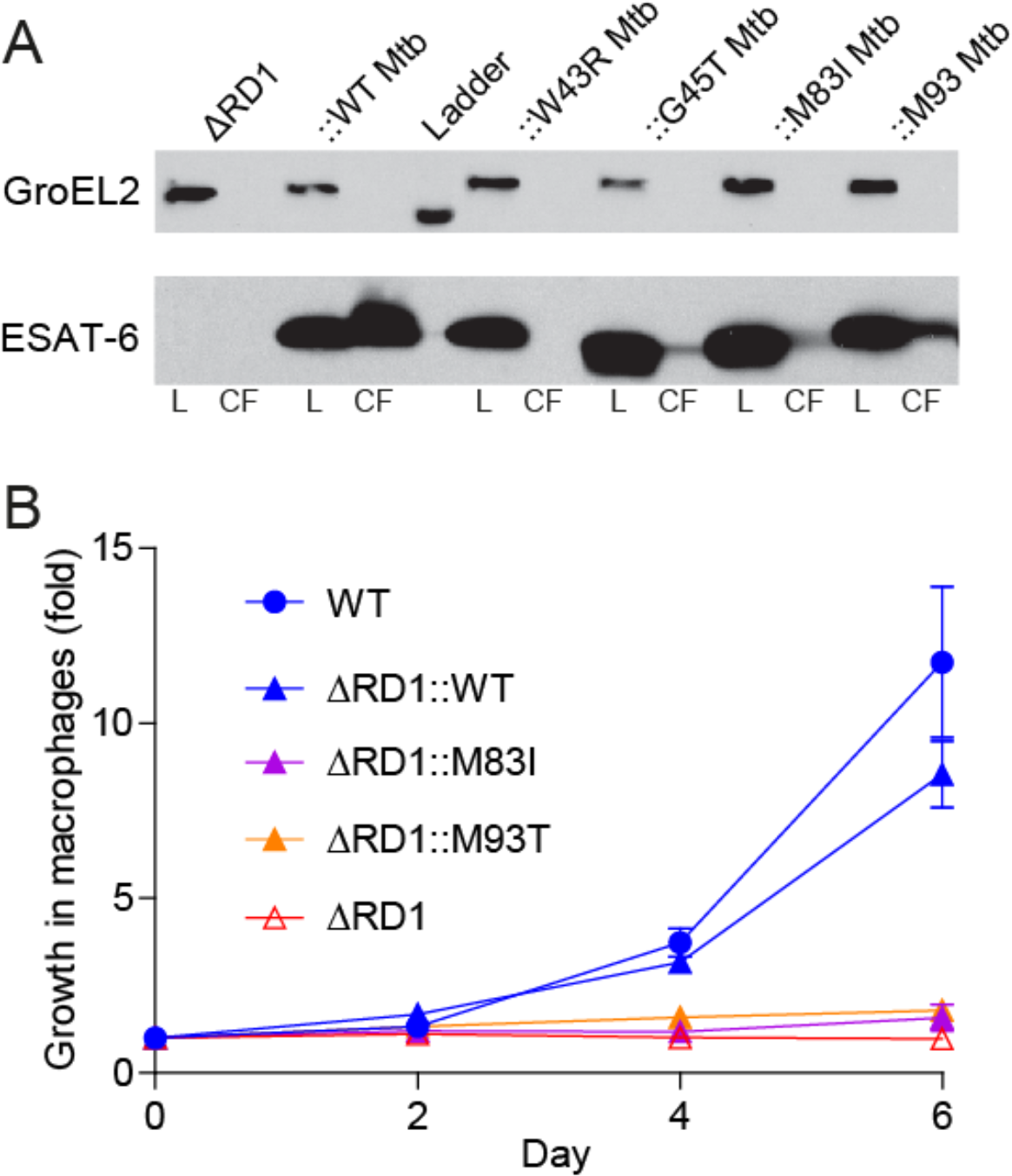
Mm ΔRD1::M83I_Mt_ and ::M93T_Mt_ mutants have reduced ESAT-6 secretion and fail to grow in macrophages. (**A**) Immunoblot of Mm lysates (L) and culture filtrates (CF) at 48 hours. Data representative of three independent experiments. (**B**) Intramacrophage growth of Mm within J774A.1 cells as measured by bacterial fluorescence. Data representative of three independent experiments.

**Figure S3.**
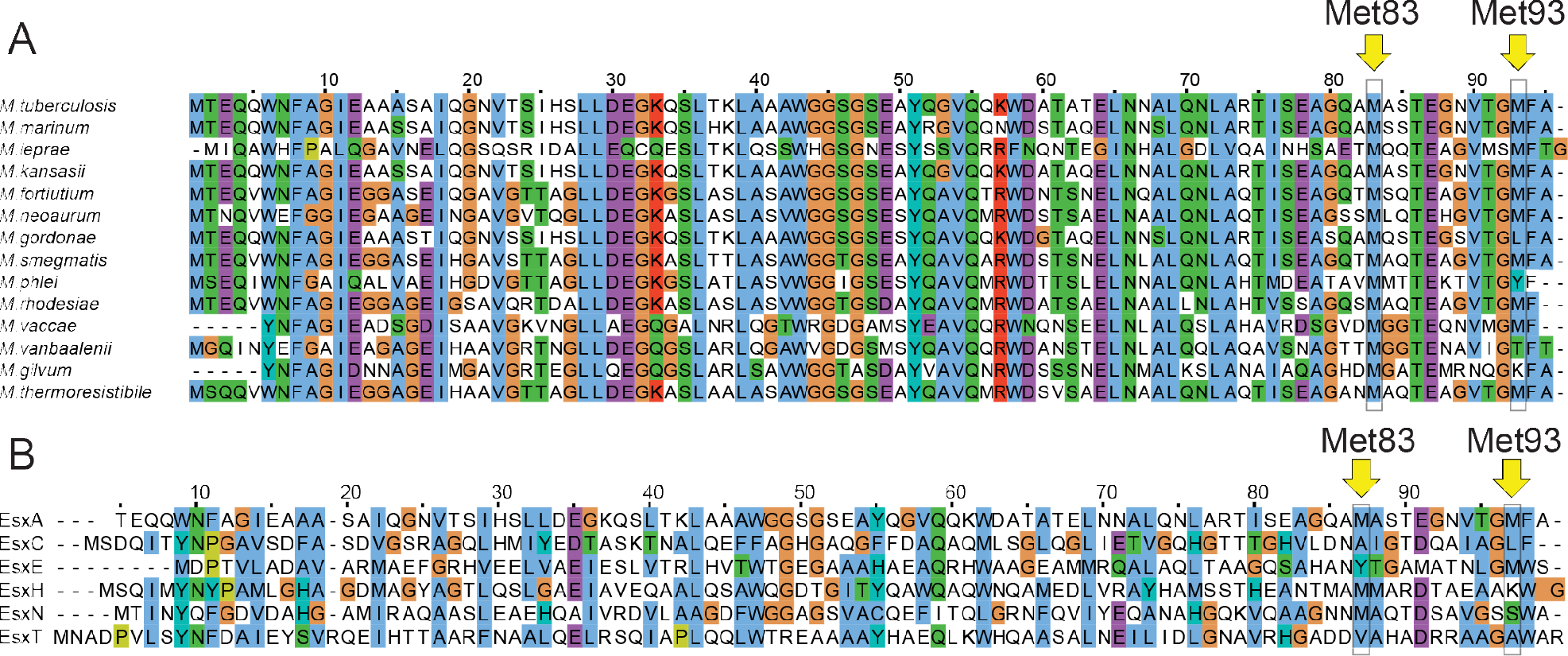
The C-terminal Met83 and Met93 residues of ESAT-6 are highly conserved. (**A**, **B**) Sequence alignment of Mtb ESAT-6 homologs (**A**) and paralogs (**B**). Yellow arrows denote residues aligned with Mtb ESAT-6 methionine 83 and 93.

### Supplementary Tables

**Table S1.**
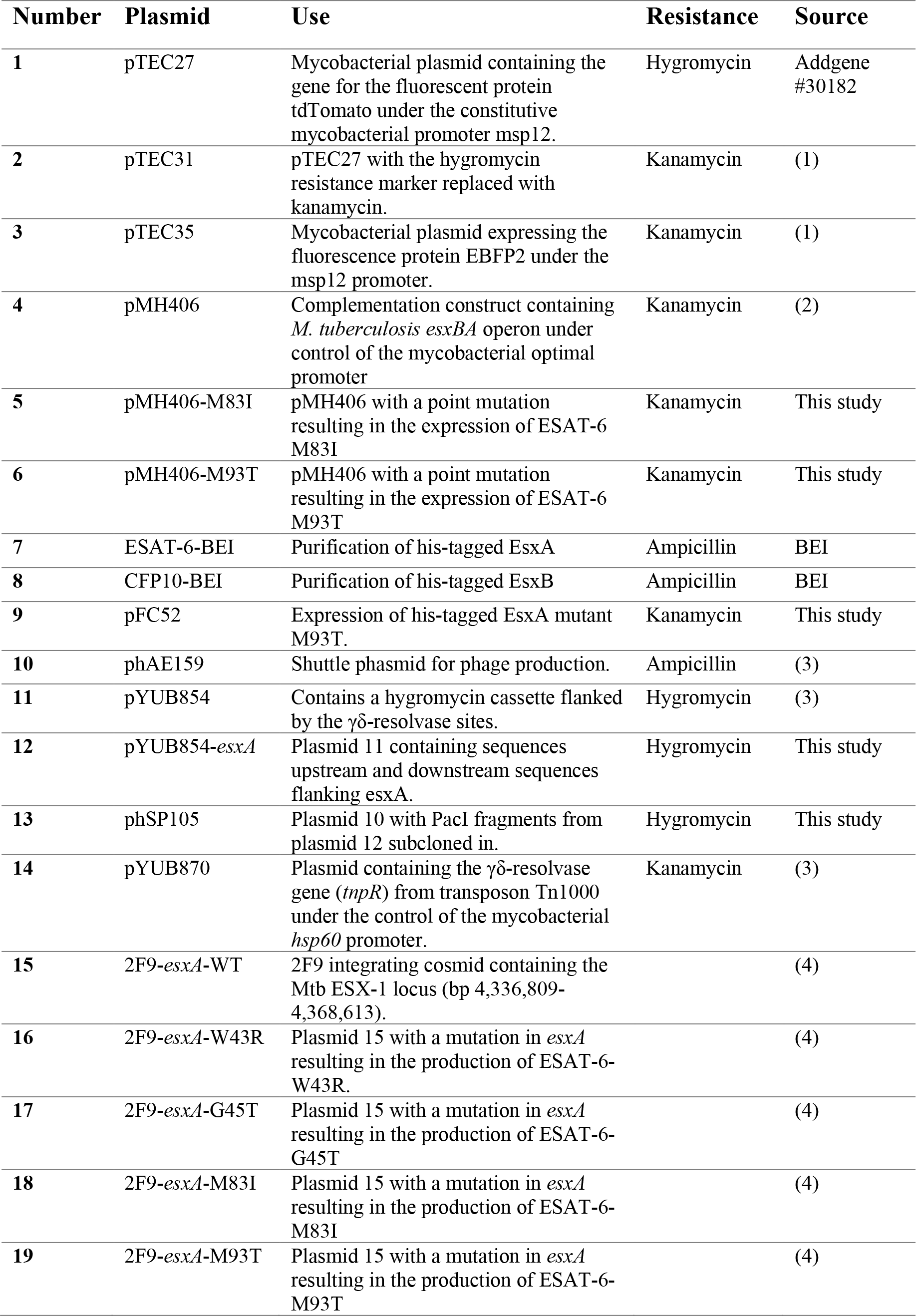
Plasmids used in this study.

**Table S2.**
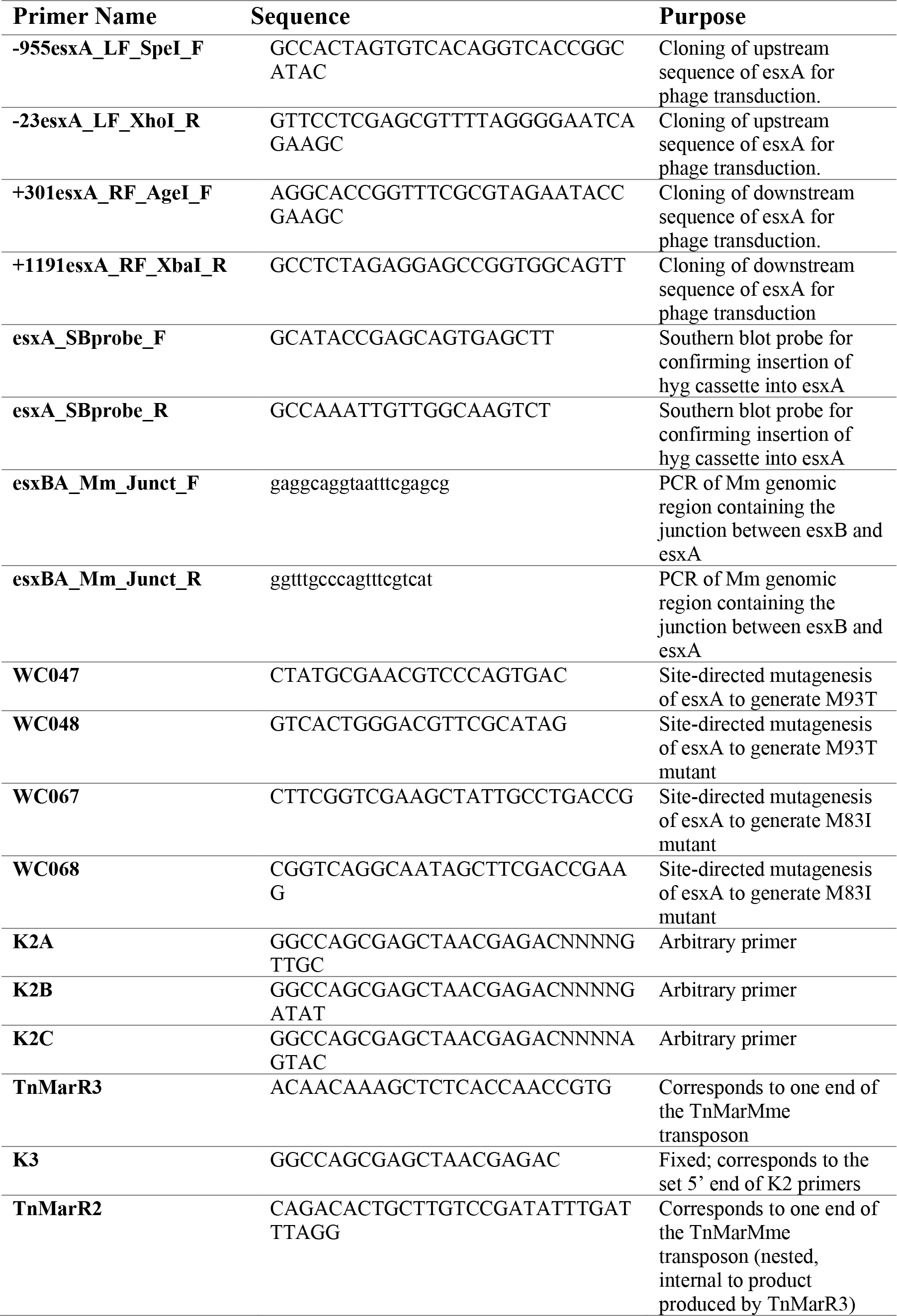
Primers used in this study.

**Table S3.**
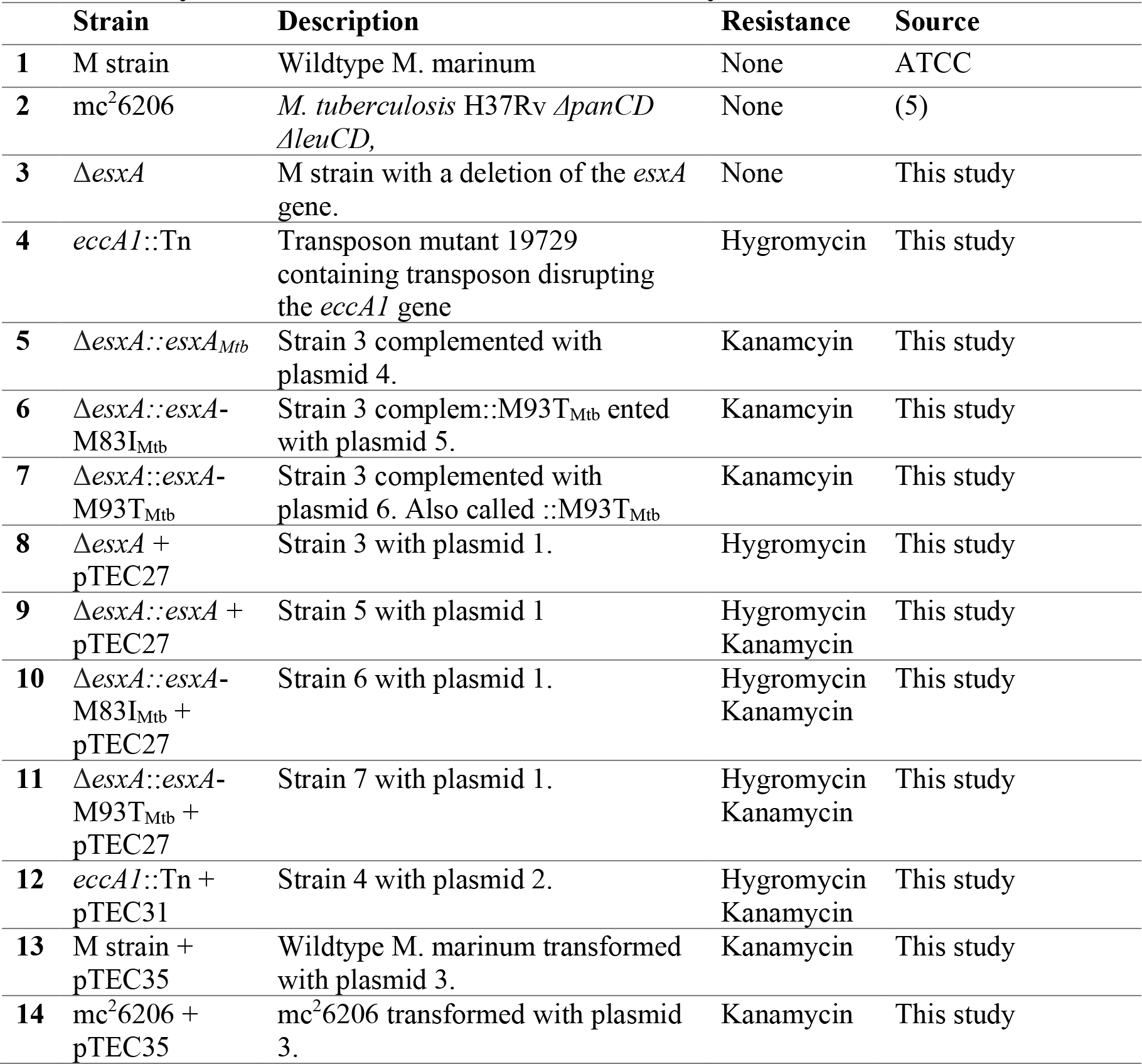
Mycobacterial strains used in this study.

